# *In silico* approach for designing of a multi-epitope based vaccine against novel Coronavirus (SARS-COV-2)

**DOI:** 10.1101/2020.03.31.017459

**Authors:** Ratnadeep Saha, Burra V L S Prasad

**Affiliations:** Department of Fisheries, Government of Tripura, India; Department of Biotechnology, K L University, Guntur, India

**Keywords:** SARS-COV-2, Immunoinformatics, Multi-epitope, Vaccine, Docking

## Abstract

A novel Coronavirus (SARS-COV-2) has now become a global pandemic. Considering the severity of infection and the associated mortalities, there is an urgent need to develop an effective preventive measure against this virus. In this study, we have designed a novel vaccine construct using computational strategies. Spike (S) glycoprotein is the major antigenic component that trigger the host immune responses. Detailed investigation of S protein with various immunoinformatics tools enabled us to identify 5 MHC I and 5 MHC II B-cell derived T-cell epitopes with VaxiJen score > 1 and IC_50_ value < 100nM. These epitopes were joined with a suitable adjuvant and appropriate linkers to form a multi-epitope based vaccine construct. Further, in silico testing of the vaccine construct for its antigenicity, allergenicity, solubility, and other physicochemical properties showed it to be safe and immunogenic. Suitable tertiary structure of the vaccine protein was generated using 3Dpro of SCRATCH suite, refined with GalaxyRefine, and validated with ProSA, PROCHECK, and ERRAT server. Finally, molecular docking studies were performed to ensure a favorable binding affinity between the vaccine construct and TLR3 receptor. The designed multi-epitope vaccine showed potential to elicit specific immune responses against the SARS-COV-2. However, further wet lab validation is necessary to confirm the actual effectiveness, safety and immunogenic potency of the vaccine construct against derived in this study.

## Introduction

A novel coronavirus 2019 (2019-nCoV), also known as severe acute respiratory syndrome coronavirus-2 (SARS-CoV-2) is a single and positive stranded RNA virus that belongs to the order Nidovirales and family Coronaviridae (Huang *et al*., 2020). The 2019-nCoV shares 79.5% and 96% of genome similarity with SARS-CoV and bat Coronavirus, respectively (Zhou *et al*., 2020; Zhu *et al*., 2020). The first incidence of cluster of pneumonia like symptoms were reported from the Wuhan city of the China in December, 2019 and the disease spread rapidly in other countries in a very short span. At last, on 11 March 2020, the Coronavirus disease 2019 (COVID-19) outbreak was officially declared as pandemic by the World Health Organization (WHO).

The outbreak of disease probably started from a single or multiple zoonotic transmission events from wet market in Wuhan, where meat and game animals were sold (Riou and Althaus, 2020). As of March 30, 2020, it has resulted in 6,93,224 confirmed cases with 33,106 deaths over 202 countries and territories (WHO Situation Report-70) [https://www.who.int/docs/default-source/coronaviruse/situation-reports/20200330-sitrep-70-covid-19.pdf?sfvrsn=7e0fe3f8_2]. Common signs of infection include fever, cough, breathing difficulties and shortness of breath. In more extreme cases, infection can cause severe acute respiratory syndrome, kidney failure and even death.

In order to control the rapidly spreading SARS-COV-2 infection, there is an urgent need to design a suitable vaccine candidate that can prevent large scale mortalities in the future. Multi-epitope based vaccines have several advantages in terms of safety, opportunity to rationally design the construct for increased efficiency, efficacy, antigenicity, and immunogenicity (Urrutia-Baca *et al*., 2019). The developmental process includes identification of virulence protein and selection of peptide segments, which can generate both cellular and humoral immune responses. In SARS-CoV spike (S) glycoprotein consists of S1 and S2 domain. S1 domains can recognize and bind to variety of receptors on the host cell surface for viral attachment. S2 domains help in fusion of host and viral membranes, allowing entry of viral genomes inside the host cells (Li, 2016). Therefore, S protein can be considered as one of the most effective target for the development vaccine against 2019-nCoV.

In the current study multiple immunoinformatics based servers and tools were used to predict T-cell epitope candidates within B-cell epitopes. Such in-silico techniques ultimately reduces the cost, effort and time compared to the traditional epitope identification approaches. Subsequently, a multi-epitope vaccine construct was designed using the most persuasive epitopes with suitable adjuvant and linkers.

## Materials and methods

### Retrieval of protein sequence

The complete amino acid sequence of surface glycoprotein or S protein of 2019-nCoV was downloaded in FASTA format from National Centre for Biotechnological Information (NCBI) database.

### Prediction of linear B-cell epitopes

Putative linear B-cell epitopes were predicted by using Artificial Neural Network (ANN) based server ABCpred (Saha and Raghava, 2006). The default threshold value of 0.51 and window length 20 was fixed for prediction.

### Identification of MHC I and MHC II epitopes within B-cell epitopes

In order to stimulate the desired and strong immune response it is critical to identify the T-cell epitopes within the B-cell epitopes. For this purpose, Propred1 and propred servers were used for identification of MHC I and MHC II binding epitopes within the pre-determined linear B-cell epitopic regions (Dar *et al*., 2019). ProPred1 is a matrix based approach uses matrices obtained from Bioinformatics and Molecular Analysis Section (BIMAS) and from the literatures (Singh and Raghava, 2003). Whereas, ProPred utilizes quantitative matrices derived from the published literature (Singh and Raghava, 2001).

MHC I epitopes were assessed with all the available 47 different alleles in Propred1 server. The option of proteasome and immunoproteasome filters was selected to improve the chances of finding accurate epitopes. Epitopes projected to be associate with at least five different MHC I alleles were retained. Whereas, MHC II epitopes were evaluated against 51 different MHC II alleles available in Propred server. Only epitopes predicted by at least ten different MHC II alleles were considered for further analysis. The predicted MHC I and MHC II binding epitopes were further subjected to VaxiJen v.2.0 server for analyzing the antigenic propensity (Doytchinova and Flower, 2007). The server was run with virus as a target field at a default threshold value of 0.4.

### Prediction of binding affinity with MHCPred

Predicted MHC I and MHC II epitopes with a VaxiJen score of >1.0 were further assessed for their binding affinity against HLA A*1101 and DRB1*0101, respectively using MHCPred version 2.0 (Guan *et al*., 2006). The epitopes with a half maximal inhibitory concentration (IC_50_) value < 100 nM were shortlisted as strong candidates for construction of multi-epitope vaccine construct associated with strong immunogenicity.

### Designing of multi-epitope based vaccine construct

For designing of a multi-epitope vaccine construct, the prioritized epitope candidates were attached with a suitable adjuvant β-defensin, and appropriate peptide linkers such as EAAAK, AAY and GPGPG.

### Evaluation antigenicity, allergenicity, solubility, and physicochemical properties

Antigenicity of the final vaccine construct was evaluated by using VaxiJen v.2.0. Whereas, Screening for allergenicity of any vaccine construct is crucial as it should not cause sensitization and allergic reaction inside the body. AllerTOP v. 2.0 (Dimitrov *et al*., 2014) and AllergenFP v.1.0 (Dimitrov *et al*., 2014) servers were used to check the allergenicity of the final vaccine construct. The solubility of vaccine construct upon expression in *Escherichia coli* was evaluated by using Protein–Sol (Hebditch *et al*., 2017). Furthermore, the various physicochemical parameters of the construct were assessed using ProtParam server (Wilkins *et al*., 1999).

### Tertiary structure prediction, refinement, and validation of vaccine protein

The three-dimensional structure of multi-epitope based vaccine was generated by using 3Dpro server of SCRATCH suite (Cheng *et al*., 2005). 3Dpro uses predicted structural features, and the Protein Data Bank (PDB) knowledge based statistical terms in the energy function. The conformational search uses a set of movements consisting on fragment substitution (using a fragment library built from the PDB), as well as random disturbance for the model. Later on the structural refinement of the modelled vaccine construct was performed through GalaxyRefine web server (Heo *et al*., 2013). This server initially reforms the side chains and executes side-chain repacking and finally overall structural relaxation by molecular dynamics simulation. Refined model was finally validated to identify any potential errors using ProSA-web, PROCHECK server and ERRAT server.

### Conformational B-cell epitope prediction of vaccine construct

DiscoTope 2.0 tool of IEDB server was used to determine conformational B-cell epitopes by using validated 3D structure of vaccine construct as an input. The server incorporates novel definition of the spatial neighborhood to sum propensity scores and half-sphere exposure as a surface measure (Kringelum *et al*., 2012).

### Molecular docking analysis and interaction studies

Molecular docking was performed in order to predict the binding affinity and interaction patterns between the vaccine construct and Toll-like receptor 3 (TLR3). The structure of TLR3 receptor was downloaded from RCSB PDB database (PDB ID: 2A0Z) and the refined 3D structure of the multi-epitope construct was used as a ligand. Finally, the binding affinity between the TLR3 receptor and vaccine construct was calculated by using ClusPro 2.0 server (Kozakov *et al*., 2017). The server uses three consecutive steps like rigid body docking, clustering of lowest energy structure, and structural refinement by energy minimization (Sayed *et al*., 2020). The best docked complex was obtained based on the lowest energy weighted score and docking efficiency. Visualization and interaction analysis of the docked complex were performed using the Chimera v1.14 and DIMPLOT program of the LigPlot^+^ v.2.1, respectively.

## Results

### Protein sequence retrieval

The complete 1273 amino acid sequence of S protein of 2019-nCoV was retrieved with an accession number **<u>YP_009724390.1</u>** to carry out the *in silico* analysis.

### Linear B-cell epitope prediction

A total 99 linear B-cell epitopes of 20 amino acid length (window length) was identified within the spike glycoprotein of 2019-nCoV. All the epitopes along with their start position and respective scores is shown in **Supplementary Table S1**.

### Identification of B-cell derived T-cell epitopes

T-cell epitope prediction comprises identification MHC I and MHC II binding epitopes, in order to activate both cytotoxic T-lymphocytes (CTL) and helper T-lymphocytes (HTL) mediated immune response. MHC I and MHC II epitopes were searched within B-cell epitopes to identify B-cell derived T-cell epitopes.

A total of 39 B-cell derived MHC I epitopes were predicted by ≥5 MHC I alleles available in Propred1. Out of these, 19 MHC I epitopes were found with above threshold VaxiJen score. A table containing MHC I binding epitopes along with number of different MHC I binding alleles and VaxiJen scores are shown in **Supplementary Table S2**. Similarly, a total of 52 B-cell derived MHC II epitopes were identified by ≥10 different MHC II alleles available in Propred server. Out of which, 26 of the MHC II epitopes were having a VaxiJen score above the fixed threshold. The MHC II epitopes with the number of encountering MHC II alleles and antigenic scores are listed in **Supplementary Table S3**. However, in order to increase the precision of selection 5 MHC I and 11 MHC II showing strong antigenicity score i.e. VaxiJen score > 1 were selected for screening of binding affinity.

### Binding affinity prediction of T-cell epitopes

Five out of five of the MHC I epitopes were found with strong binding affinity against HLA A*1101 allele, and five out of eleven MHC II epitopes were having favorable binding affinity against DRB1*0101 in human. Finally B-cell derived T-cell epitopes were selected based on our set of criteria like the total number of binding alleles (≥5 for MHC I, and ≥10 for MHC II), VaxiJen score > 1.0 and binding affinity (IC_50_ < 100 nM) is shown in **Table 1 and 2**.

**Table 1.** Selected five of the B-cell derived MHC I binding peptides based on our set of pre-defined criteria i. Total number of different binding alleles must be ≥5, ii. VaxiJen score >1.0, and iii. Binding affinity against HLA A*1101 (IC50 < 100 nM).

| Sl. No. | Start Position | Sequence | No. of MHC I binding alleles | VaxiJen 2.0 Score | MHCPred HLA A*1101 $IC_{50}$ Value (nM) |
| --- | --- | --- | --- | --- | --- |
| 1. | 109 | TLDSKTQSL | 12 | 1.0685 | 82.6 |
| 2. | 181 | GKQGNFKNL | 8 | 1.0607 | 22.03 |
| 3. | 379 | CYGVSP TKL | 6 | 1.4263 | 79.62 |
| 4. | 417 | KIADYNYKL | 13 | 1.6639 | 52.97 |
| 5. | 510 | VVVL SFELL | 14 | 1.0909 | 16.52 |

**Table 2.** Selected five out of eleven B-cell derived MHC II binding peptides (shown in bold) based on our set of pre-defined criteria i. Total number of different binding alleles must be ≥10, ii. VaxiJen score >1.0, and iii. Binding affinity against DRB1*0101 (IC50 < 100 nM).

| Sl. No. | Start Position | Sequence | No. of MHC I binding alleles | VaxiJen 2.0 Score | MHCPred DRB1*0101 $IC_{50}$ Value (nM) |
| --- | --- | --- | --- | --- | --- |
| **1.** | **231** | **IGINITRFQ** | **35** | **1.3386** | **98.4** |
| **2.** | **495** | **YGFQPTNGV** | **12** | **1.0509** | **37.33** |
| 3. | 511 | VVLSFELLH | 11 | 1.409 | 171.79 |
| **4.** | **512** | **VLSFELLHA** | **20** | **1.0776** | **41.5** |
| 5. | 534 | VKNKCVNFN | 16 | 2.053 | --- |
| **6.** | **894** | **LQIPFAMQM** | **17** | **1.068** | **6.43** |
| 7. | 1060 | VVFLHV TYV | 44 | 1.5122 | 883.08 |
| 8. | 1061 | VFLHV TYVP | 13 | 1.2346 | 488.65 |
| 9. | 1128 | VVIGIVNNT | 21 | 1.3063 | 203.24 |
| 10. | 1172 | INASVVNIQ | 18 | 1.1445 | 289.07 |
| **11.** | **1225** | **IAIVMVTIM** | **15** | **1.1339** | **47.75** |

### Multi-epitope vaccine design

For designing of a 183 amino acid long multi-epitope based vaccine construct, prioritized 5 MHC I and 5 MHC II binding epitopes were merged together by using AAY and GPGPG linkers, respectively. Further, N-terminal end of the first MHC I epitope was joined with the β-defensin adjuvant by using EAAAK linker. Whereas, the C-terminal end of the last MHC II binding epitope was linked with 6x-His tag. The sequence of the designed vaccine construct is given below:

GIINTLQKYYCRVRGGRCAVLSCLPKEEQIGKCSTRGRKCCRRKK**EAAAK**TLDSKTQSLAAYGKQGNFKNL**AA Y**CYGVSPTKLAAYKIADYNYKL**AAY**VVVLSFELL**GPGPG**IGINITRFQ**GPGPG**YGFQPTNGV**GPGPG**VLSFELL HA**GPGPG**LQIPFAMQM**GPGPG**IAIVMVTIM**HHHHHH**

### Prediction of antigenicity, allergenicity, solubility, physicochemical property

The designed vaccine construct was predicted as antigenic, non-allergen and soluble in nature. Various physicochemical properties of the vaccine constructs are listed in **Table 3**.

**Table 3.** Antigenicity, allergenicity, solubility, and physicochemical property assessments of the primary sequence of multi-epitope based vaccine construct.

| Sl. No. | Features | Assessment |
| --- | --- | --- |
| 1. | Antigenicity | 0.6243 (Probable ANTIGEN) |
| 2. | Allergenicity | Probable non-allergen (AllerTOP v.2.0)<br>Probable non-allergen (AllergenFP v.1.0) |
| 3. | Solubility | 0.578 (Soluble) |
| 4. | Number of amino acids | 183 |
| 5. | Molecular weight | 19572.85 Dalton |
| 6. | Theoretical Isoelectric point (pI) | 9.66 |
| 7. | Total number of atoms | 2761 |
| 8. | Formula | C<sub>879</sub>H<sub>1386</sub>N<sub>248</sub>O<sub>237</sub>S<sub>11</sub> |
| 9. | Estimated half-life | 30 hours (mammalian reticulocytes, in vitro)<br>>20 hours (yeast, in vivo)<br>>10 hours (Escherichia coli, in vivo) |
| 10. | Instability index | 25.48 (Stable) |
| 11. | Aliphatic index | 80.00 |
| 12. | Grand average of hydropathicity (GRAVY) | -0.111 |

### Tertiary structure prediction, refinement and validation of vaccine protein

Due to unavailability of good structural templates for homology modeling in the PDB database, 3Dpro was suitable to build the model of vaccine structure. Refined 3D model by GalaxyRefine is shown in **Fig. 1A**.

**Fig 1.**
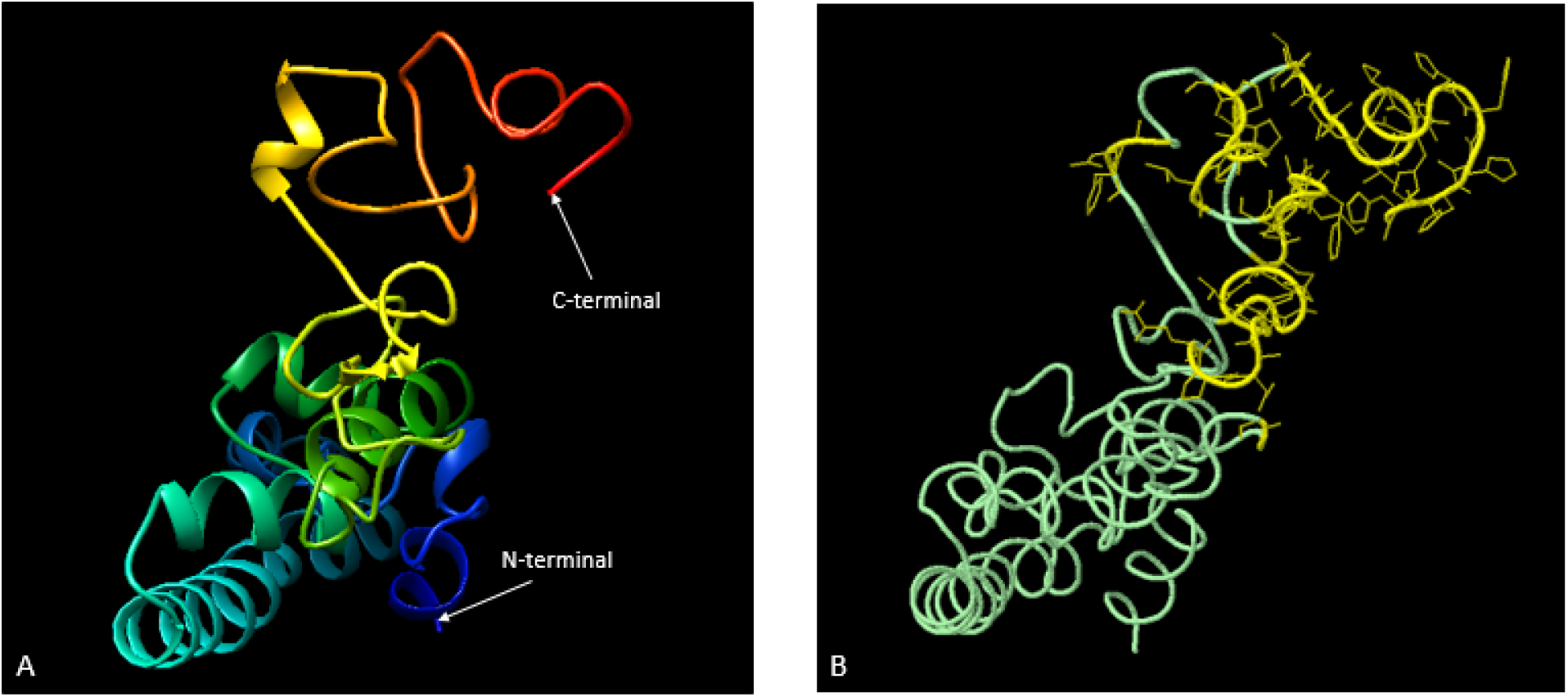
3D modelling and conformational B-cell epitopes of the multi-epitope based vaccine construct: A. Refined tertiary structure of the vaccine protein; B. Discotope 2.0 prediction of conformational epitopes (in yellow).

Ramachandran plot by PROCHECK confirms 90.8, 7.7, 0.0, and 1.4% of the residues were in most favoured regions, additional allowed regions, generously allowed regions and disallowed regions, respectively **(Fig. 2A)**. ProSA Z-score of the refined model was found to be -4.42, which falls within the vicinity of experimental structures **(Fig. 2B)**. ProSA also showed a valid local model quality by plotting energies as a function of amino acids present in protein structure **(Fig. 2C)**. Overall quality score by ERRAT was projected as 85.714, which further supports the refined structure as the high quality model **(Fig. 2D)**.

**Fig 2.**
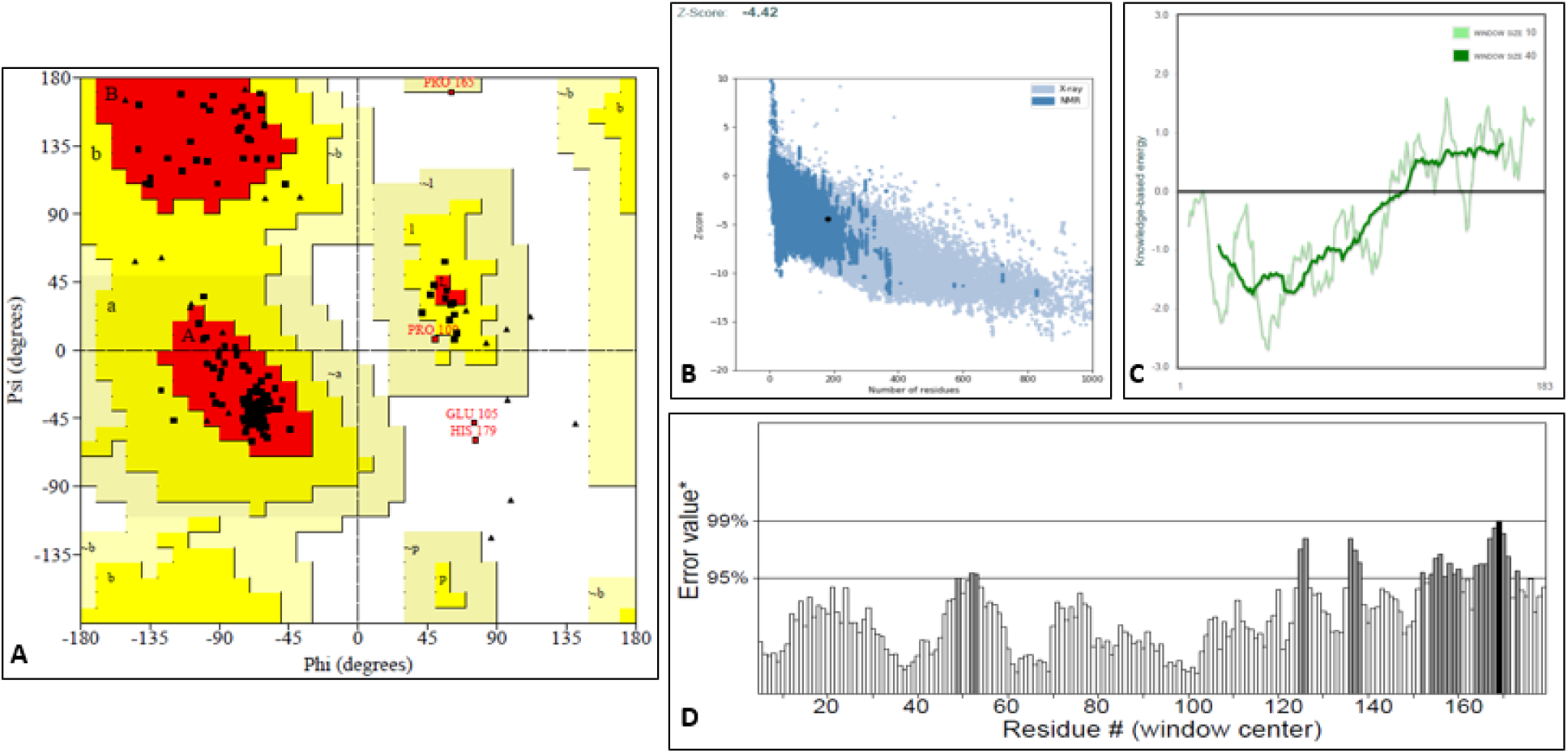
Structural validation of the refined modeled vaccine: A. Ramachandran plot generated using PROCHECK. The areas showing different colors i.e. red, yellow and light yellow represents most favored regions (90.8%), additional allowed regions (7.7%), and disallowed regions (1.4%) respectively. B. ProSA Z-score (Overall model quality); C. ProSA graphical plot (Local model quality); D. The ERRAT plot.

### Conformational B-cell epitope prediction

A total of 43 residues out of 183 amino acids of the vaccine construct were predicted as discontinuous B-cell epitopes at an above DiscoTope score threshold -3.70 **(Fig. 1B)**. The predicted conformational epitope sequences are given below:

**G**^**112**^**-QPTNGVGPGP**^**130-139**^**-FE**^**144-145**^**-GPGPGL**^**150-155**^**-IPFAMQM**^**157-163**^**-GPG**^**166-168**^**-AIVMVTIMHHHHHH**^**170-183**^ Detailed prediction of contact number, propensity score, and Discotope score against each residues are shown in **Supplementary Table S4**.

### Molecular docking of vaccine with TLR3 receptor

Total 30 models were generated through ClusPro 2.0 server showing the interaction between the refined vaccine construct and TLR3 receptor. A table containing all the energy scores of docked complexes is shown in **Supplementary Table S5**. Among all the docked models, the model number 4 was selected as the best docked complex due to its lowest energy score i.e. -1119.2, indicating highest binding affinity, showed in **Fig. 3 (A-B)**.

**Fig 3.**
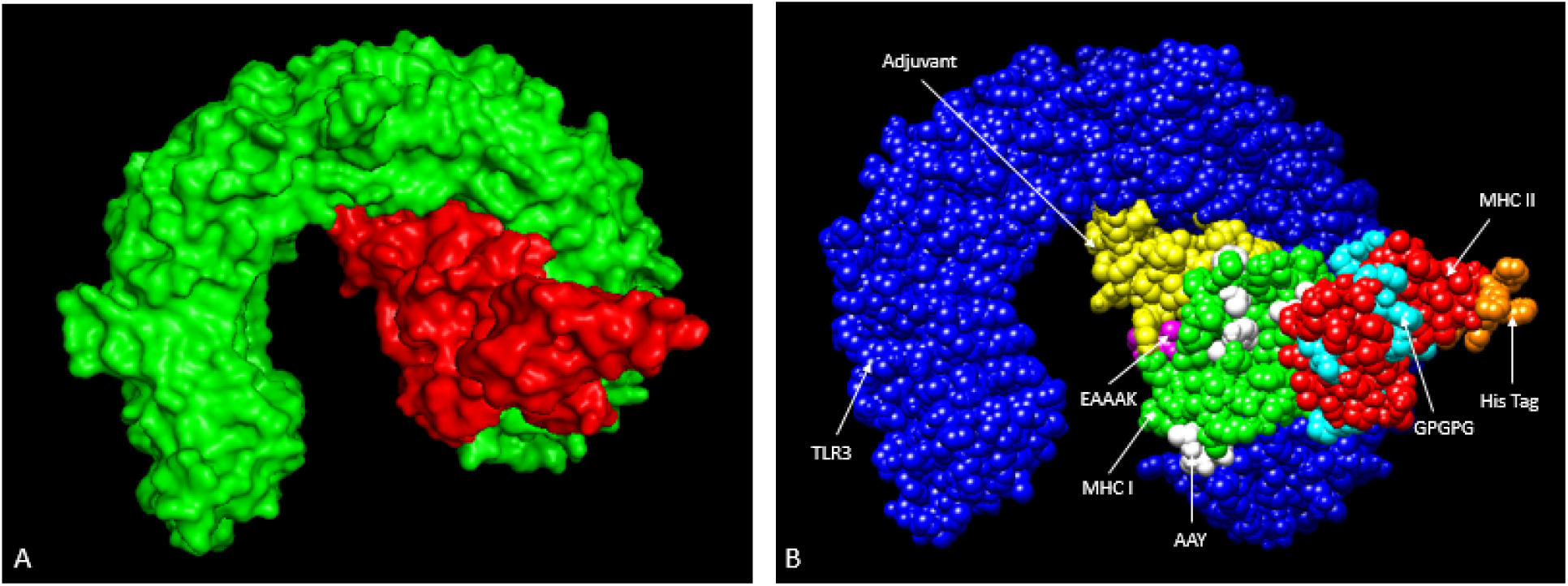
Docked complex of the modeled vaccine construct and TLR3 receptor: A. Complex showing the surface interaction between vaccine component (green) and TLR3 (red); B. Docked complex in sphere view (TLR3 receptor: Blue, β-defensin adjuvant: Yellow, EAAAK linker: Magenta, AAY linker: White, GPGPG linker: Cyan, MHC I epitopes: Green, MHC II epitopes: Red, His tag: Orange).

The intra-molecular interactions including hydrogen bonds and hydrophobic interactions are represented in **Fig 4 (A-D)**. The DIMPLOT analysis showed that nineteen residues of the multi-epitope vaccine construct were involved in formation of 31 nos. of hydrogen bonds with TLR3. The length of hydrogen bonds was ranged from 2.55 to 3.04 Å.

**Fig 4.**
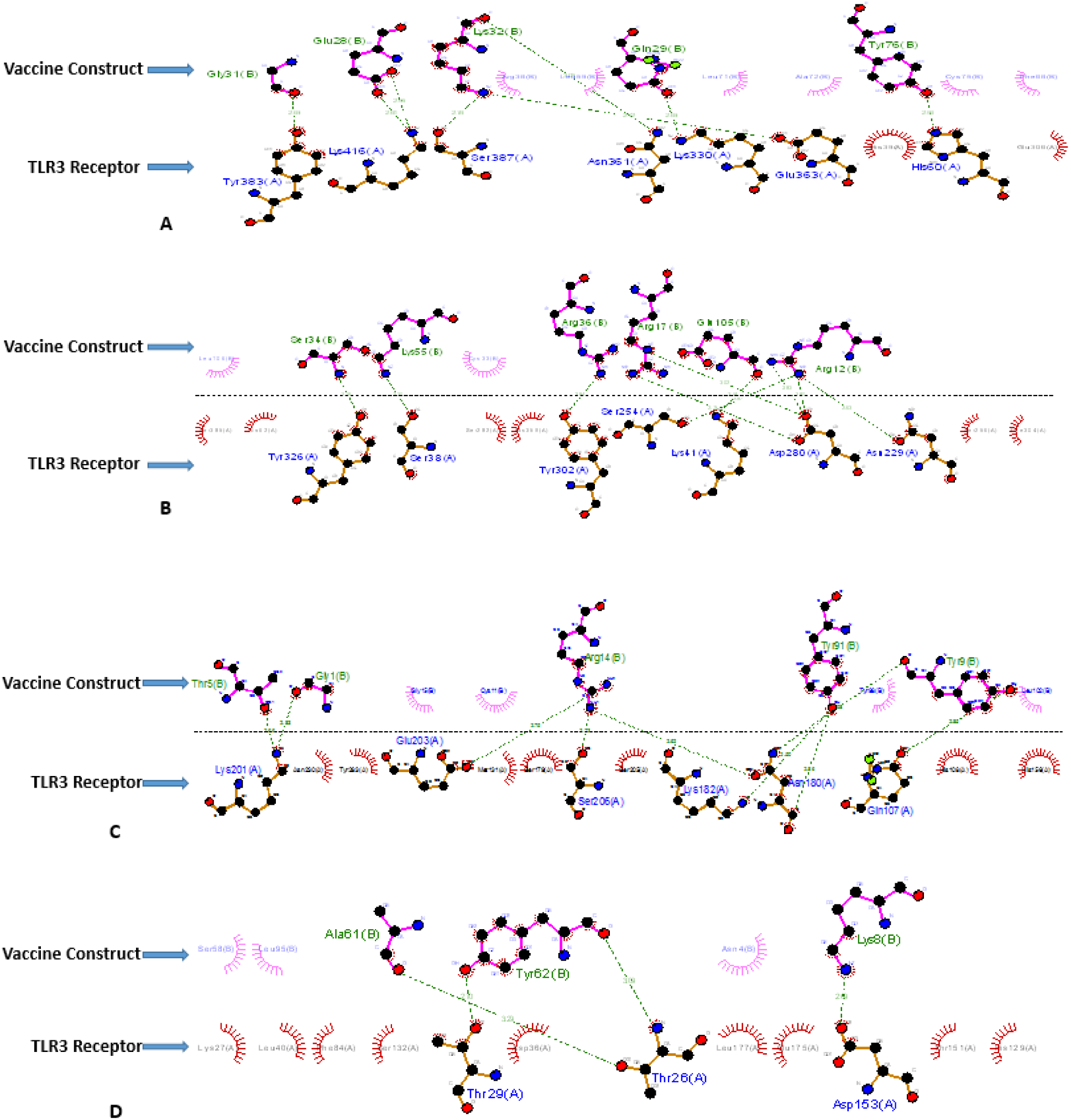
2D interaction studies by using DIMPLOT: (A-D) Vaccine construct (Chain B) showing hydrogen bonding (green dotted lines) and hydrophobic interactions (arcs with lines) with TLR3 receptor (Chain A).

## Discussion

The 2019-nCoV has become pandemic, showing no sign of abatement. The virus is highly contagious in nature, causing respiratory distress that can led to eventual death in susceptible individuals. Researchers across the world are fraught with the challenge of finding means for halting the spread of this virus. Historically, vaccination has proved to be an effective method of protecting large human population against viral diseases. Comparison to traditional vaccines like killed, attenuated or live vaccines, epitope based vaccines which contain specifically targeted immunogenic component of the pathogen responsible for causing diseases can be designed more rationally (Pourseif *et al*., 2019).

In the present study, we focused on the identification of potential B-cell derived T-cell epitopes in order to generate a potential vaccine construct which can induce both humoral and cellular immune responses simultaneously. Several authors reported that in order to produce a desired immune response, the epitopes must be accessible to both MHC I and MHC II molecules along with the B-cell (Patra *et al*., 2019). Out of several viral proteins, S protein based vaccines and antiviral therapies have been reported to be effective against the earlier encounters of SARS-CoV and MERS-CoV infections (Du *et al*., 2009; Schindewolf and Menachery, 2019). With the help of immunoinformatics approaches, we were able to identify 5 MHC I and 5 MHC II epitopes with high antigenic potential and strong binding affinity within the S protein of 2019-nCoV. Binding affinity of the selected epitopes was predicted against HLA A*1101 and DRB1*0101 alleles, as these are the most common MHC class I and MHC class II alleles, respectively in human population. Bhattacharya *et al*., (2020) also predicted and designed an epitope based peptide vaccine against 2019-nCoV. However, our study revealed eight out of ten completely different sets of epitopes with better Vaxijen scores i.e. > 1, which have not been reported earlier.

For designing of a multi-epitope based vaccine construct, predicted epitopes were joined together using different linkers for adequate separation of the epitopes. A suitable adjuvant was added at the N-terminus end to boost the immunogenicity within the human body and 6x-His tag was added to the C-terminus end for identification and purification purpose. The vaccine construct was predicted as probable antigen, non-allergen and soluble in nature. So, the designed multi-epitope vaccine have the potential to produce more effective, specific, robust, and durable immune response without causing any adverse effect in humans.

Protparam analysis of the multi-epitope construct reveals the pI (theoretical isoelectric point) value 9.66, which means the vaccine protein is basic in nature and most stable at this pH range. Further, the aliphatic index 80.00 indicates thermostable nature of the construct at various temperature and the instability index 25.48 (<40) shows that construct will remain stable after expression. The Grand average of hydropathicity (GRAVY) value was computed to be negative (−0.111), reveals that vaccine is hydrophilic in nature and likely to interact with other protein molecules.

Further, the generated refined tertiary structure of vaccine construct was found valid for identification of conformational epitopes and docking experiments. ProSA Z score showed the overall quality score of the model protein, which fallen within the range characteristics of the native protein. The PROCHECK Ramachandran plot used to find out energetically allowed and disallowed psi (ψ) and phi (□) dihedral angles of amino acids, which was calculated based on van der Waal radius of the side chain. Our modeled vaccine construct showed only less than 1.5% of the residues were present in disallowed regions of Ramachandran plot, indicating negligible amount of steric clashes between the side chain atoms and main chain atoms. Again ERRAT server was used to find out the pattern of non-bonded atomic interactions. The overall quality factor (ERRAT score) was >50 i.e. 85.714, indicates the high quality model.

Further Discotope 2.0 identified 43 residues as conformational B-cell epitopes within the vaccine construct. These epitopes can comes in a close contact to form a three dimensional conformation due to protein folding, which can be identified by B-cells. Molecular docking study was carried out to analyze the interaction between the vaccine construct and TLR3 receptor. The lowest stabilized energy score by Cluspro 2.0 indicated strong and favorable interaction of our multi-epitope vaccine construct with innate immune receptor, which can ultimately activate TLR and augment the immune response against 2019-nCoV.

## Conclusion

2019-nCoV is a new virus which become a serious Public Health Emergency of Global Concern. Current study followed a reverse vaccinology approach to identify high ranked epitopes using an immunoinformatics approach, to formulate a novel multi-epitope based vaccine construct to prevent this disastrous outbreak. The designed vaccine construct has suitable structural, physicochemical and immunological properties which can strongly stimulate both humoral and cellular immune responses in humans. However, the proposed vaccine construct must be validated through *in-vitro* and *in-vivo* bioassays to prove its safety, efficacy, and immunogenicity against COVID-19.

## Supplementary data

**Supplementary Table S1**. List of all predicted 20-mer linear B-cell epitopes by ABCpred along with their sequence, start position and score

**Supplementary Table S2**. List of identified B-cell derived 9-mer MHC I epitopes predicted by Propred1 along with their VaxiJen score

**Supplementary Table S3**. List of identified B-cell derived 9-mer MHC II epitopes predicted by Propred along with their VaxiJen score

**Supplementary Table S4**. Predicted conformational B-cell epitope residues of multi-epitope vaccine construct using DiscoTope 2.0

**Supplementary Table S5**. Predicted top 30 energy scores between vaccine-receptor docked complexes

## References

1. Bhattacharya M, Sharma AR, Patra P, Ghosh, P., Sharma, G., Patra, B.C., Lee, S.S., Chakraborty, C., 2020. Development of epitope-based peptide vaccine against novel coronavirus 2019 (SARS-COV-2): Immunoinformatics approach. J. Med. Virol. doi: 10.1002/jmv.25736

2. Cheng, J., Randall, A.Z., Sweredoski, M.J., Baldi, P., 2005. SCRATCH: A protein structure and structural feature prediction server. Nucleic Acids Res. 33, W72–W76. doi: 10.1093/nar/gki396

3. Dar, H. A., Zaheer, T., Shehroz, M., Ullah, N., Naz, K., Muhammad, S. A., Zhang, T., Ali, A., 2019. Immunoinformatics-Aided Design and Evaluation of a Potential Multi-Epitope Vaccine against *Klebsiella Pneumoniae*. Vaccines (Basel), 7, doi: 10.3390/vaccines7030088.

4. Dimitrov, I., Bangov, I., Flower, D. R., Doytchinova, I., 2014. AllerTOP v.2 - A server for *in silico* prediction of allergens, J. Mol. Model. 20, 2278–2284. doi:10.1007/s00894-014-2278-5.

5. Dimitrov, I., Naneva, L., Doytchinova, I., Bangov, I., 2014. AllergenFP: Allergenicity prediction by descriptor fingerprints. Bioinformatics, 30, 846–851. doi:10.1093/bioinformatics/btt619.

6. Doytchinova, I.A., Flower, D.R., 2007. VaxiJen: a server for prediction of protective antigens, tumour antigens and subunit vaccines. BMC Bioinformatics, 8, 4. doi: 10.1186/1471-2105-8-4

7. Du, L., He, Y., Zhou, Y., Liu, S., Zheng, B.J., Jiang, S., 2009. The spike protein of SARS-CoV--a target for vaccine and therapeutic development. Nat. Rev. Microbiol. 7, 226–236. doi: 10.1038/nrmicro2090

8. Guan, P., Hattotuwagama, C.K., Doytchinova, I.A., Flower, D.R., 2006. MHCPred 2.0: an updated quantitative T-cell epitope prediction server. Appl. Bioinformatics, 5, 55–61. doi: 10.2165/00822942-200605010-00008

9. Hebditch, M., Carballo-Amador, M.A., Charonis, S., Curtis, R., Warwicker, J., 2017. Protein-Sol: a web tool for predicting protein solubility from sequence. Bioinformatics, 33, 3098–3100. doi: 10.1093/bioinformatics/btx345

10. Heo, L., Park, H., and Seok, C., 2013. GalaxyRefine: Protein structure refinement driven by side-chain repacking. Nucleic Acids Res., 41, W384–W388. doi: 10.1093/nar/gkt458

11. Huang, C., Wang, Y., Li, X., et al., 2020. Clinical features of patients infected with 2019 novel coronavirus in Wuhan, China. Lancet. 395, 497–506. doi: 10.1016/S0140-6736(20)30183-5

12. Kozakov, D., Hall, D.R., Xia, B., Porter, K. A., Padhorny, D., Yueh, C., Beglov, D., Vajda, S., 2017. The ClusPro web server for protein–protein docking. Nat. Protoc. 12, 255–278. doi:10.1038/nprot.2016.169

13. Kringelum, J.V., Lundegaard, C., Lund, O., Nielsen, M., 2012. Reliable B Cell Epitope Predictions: Impacts of Method Development and Improved Benchmarking. PLoS Comput. Biol. 8, e1002829. doi: 10.1371/journal.pcbi.1002829

14. Li, F., 2016. Structure, Function, and Evolution of Coronavirus Spike Proteins. Annu. Rev. Virol. 3, 237–261. doi: 10.1146/annurev-virology-110615-042301

15. Patra, P., Mondal, N., Patra, B.C., Bhattacharya, M., 2019. Epitope-Based Vaccine Designing of *Nocardia asteroides* Targeting the Virulence Factor Mce-Family Protein by Immunoinformatics Approach. Int. J. Pept. Res. Ther. doi: 10.1007/s10989-019-09921-4

16. Pourseif, M.M., Yousefpour, M., Aminianfar, M., Moghaddam, G., Nematollahi, A., 2019. A multi-method and structure-based *in silico* vaccine designing against *Echinococcus granulosus* through investigating enolase protein. Bioimpacts, 9, 131–144. doi: 10.15171/bi.2019.18

17. Riou, J., Althaus, C.L., 2020. Pattern of early human-to-human transmission of Wuhan 2019 novel coronavirus (2019-nCoV), December 2019 to January 2020. Euro. Surveill. 25, 20200220c. doi: 10.2807/1560-7917.ES.2020.25.4.2000058

18. Saha, S., Raghava, G.P.S., 2006. Prediction of Continuous B-cell Epitopes in an Antigen Using Recurrent Neural Network. Proteins, 65, 40–48. doi: 10.1002/prot.21078

19. Sayed, S.B., Nain, Z., Khan, M.S.A., Abdulla, F., Tasmin, R., Adhikari, U.K., 2020. Exploring Lassa Virus Proteome to Design a Multi-epitope Vaccine Through Immunoinformatics and Immune Simulation Analyses. Int. J. Pept. Res. Ther. doi: 10.1007/s10989-019-10003-8

20. Schindewolf, C., Menachery, V.D., 2019. Middle East Respiratory Syndrome Vaccine Candidates: Cautious Optimism. Viruses, 11, doi: 10.3390/v11010074

21. Singh, H., Raghava, G.P.S., 2001. ProPred: Prediction of HLA-DR binding sites. Bioinformatics, 17, 1236–1237. doi: 10.1093/bioinformatics/17.12.1236

22. Singh, H., Raghava, G.P.S., 2003. ProPred1: Prediction of promiscuous MHC class-I binding sites. Bioinformatics, 19, 1009–1014. doi: 10.1093/bioinformatics/btg108

23. Urrutia-Baca, V.H., Gomez-flores, R., Garza-Ramos, M.A.D.L., Tamez-guerra, P., Lucio-sauceda, D.G., Rodríguez-padilla, M. C., 2019. Immunoinformatics Approach to Design a Novel Epitope-Based Oral Vaccine Against *Helicobacter pylori*. J. Comput. Biol. 26, 1177–1190. doi: 10.1089/cmb.2019.0062

24. Wilkins, M.R., Gasteiger, E., Bairoch, A., Sanchez, J.C., Williams, K.L., Appel, R.D., Hochstrasser, D. F., 1999. Protein identification and analysis tools in the ExPASy server. Methods Mol. Biol. 112, 531–552. doi: 10.1385/1-59259-584-7:531

25. Zhou, P., Yang, X-L., Wang, X-G., et al., 2020. A pneumonia outbreak associated with a new coronavirus of probable bat origin. Nature, 579, 270–273. doi: 10.1038/s41586-020-2012-7

26. Zhu, N., Zhang, D., Wang, W., et al., 2020. A novel coronavirus from patients with pneumonia in China, 2019. N. Engl. J. Med. 382, 727–733. doi: 10.1056/NEJMoa2001017

